# Mouse embryonic stem cells switch migratory behaviour during early differentiation

**DOI:** 10.1101/2020.12.07.415307

**Authors:** Irene M. Aspalter, Wolfram Pönisch, Kevin J. Chalut, Ewa K. Paluch

## Abstract

Development relies on a series of precisely orchestrated cell fate changes. While studies of fate transitions often focus on changes in gene regulatory networks, most transitions are also associated with changes in cell shape and cell behaviour. Here, we investigate changes in migratory behaviour in mouse embryonic stem (ES) cells during their first developmental fate transition, exit from ES cell state. We show that naïve pluripotent ES cells cannot efficiently migrate on 2-dimensional substrates but are able to migrate in an amoeboid fashion when placed in confinement. Exit from ES cell state, typically characterised by enhanced cell spreading, is associated with decreased migration in confinement and acquisition of mesenchymal-like migration on 2D substrates. Interestingly, confined, amoeboid-like migration of ES cells strongly depends on Myosin IIA, but not Myosin IIB. In contrast mesenchymal-like migration of cells exiting the ES cell state does not depend on Myosin motor activity but relies on the activity of the Arp2/3 complex. Together, our data suggest that during early differentiation, cells undergo a switch in the regulation of the actin cytoskeleton, leading to a transition from amoeboid-to mesenchymal-like migration.

**Summary statement:** Naïve mouse embryonic stem cells display amoeboid-like migration in confinement, but switch to mesenchymal-like migration as they exit the ES cell state.

## Introduction

Transitions in cellular states are of fundamental importance for embryonic development, tissue homeostasis, and disease progression. Cell state transitions have been thoroughly investigated from the perspective of the underlying changes in genetic regulatory networks (Rue and Martinez Arias, 2015, Pastushenko and Blanpain, 2019, Kalkan et al., 2017). However, cell state transitions are also generally associated with alterations in the regulation and execution of key cellular processes, such as cell division and migration. Such transitions in cell behaviour have been extensively studied in cancer, where changes in division and migration are key features of disease progression (Wolf and Friedl, 2006, Matthews et al., 2020). In contrast, the cellular changes associated with state transitions in development, in particular in mammals, have received less attention. This is in part due to the difficulty of investigating individual cell behaviour in mammalian embryos. Yet, recent studies highlight the importance of changes in cellular properties, such as cell surface tension, adhesiveness, and extracellular matrix interactions, during early mouse development (Kyprianou et al., 2020, Samarage et al., 2015, Maitre et al., 2015).

Mouse embryonic stem (ES) cells, isolated from the inner cell mass of the early blastocyst, are a good model system for studying cell state transitions of early mouse development (Kalkan et al., 2017). ES cells can be maintained in culture in a naïve ground state of pluripotency, and their gene expression patterns are comparable to epiblast cells *in vivo* (Nichols and Smith, 2009, Boroviak et al., 2014). When microinjected into another blastocyst, ES cells can contribute to all lineages showing full pluripotent capacity (Gardner, 1998). Changing the culture conditions *in vitro* causes ES cells to exit naïve pluripotency (the ES cell state) and begin the process of differentiation (Kalkan et al., 2017, Kalkan and Smith, 2014). Interestingly, exit from the ES cell state is associated with cellular shape changes from round morphologies in colonies of ES cells to spread shapes displayed by cells exiting naïve pluripotency (Chalut and Paluch, 2016, De Belly et al., 2020).

The shape change displayed by ES cells during early differentiation is reminiscent of the morphological changes characterizing amoeboid-to-mesenchymal transitions in migrating cells (Yamada and Sixt, 2019). Cells migrating in 3-dimensional (3D) environments can display a variety of migration modes, including mesenchymal and amoeboid, and many cell types are able to transition between modes depending on internal and external conditions (Yamada and Sixt, 2019, Te Boekhorst et al., 2016, Bodor et al., 2020). Amoeboid cells display rounded morphologies, often form bleb-like protrusions, and do not require focal adhesions for migration when placed in confinement (reviewed in (Paluch et al., 2016)). Some amoeboid cells, such as Dictyostelium, can migrate over 2D substrates, but others appear to only become motile in confinement, presumably because their levels of substrate adhesiveness are insufficient to effectively propel cells in 2D (Lammermann and Sixt, 2009, Liu et al., 2015). In contrast, mesenchymal cells display spread morphologies with protrusive lamellipodia, and can migrate on 2D substrates, using focal adhesions to exert propelling forces (Te Boekhorst et al., 2016).

Both migration modes rely on the actomyosin cytoskeleton for propelling force generation (Callan-Jones and Voituriez, 2016) through non-muscle Myosins (hereafter Myosins) (Agarwal and Zaidel-Bar, 2019), but differ in requirement for other cytoskeletal components (Bergert et al., 2012, Lehtimaki et al., 2017, Paluch et al., 2016). A key cytoskeletal regulator of mesenchymal migration is generally thought to be the Arp2/3 complex, as it nucleates the lamellipodial actin network extending the leading edge of the cell (Swaney and Li, 2016). In contrast, Arp2/3 activity is not required for the amoeboid migration of confined cells (Bergert et al., 2012), where both protrusion formation and propelling force generation primarily rely on actomyosin contractility. During amoeboid migration, Myosin-generated contractile forces drive the formation of blebs at the front of the cell, and the generation of actomyosin retrograde flows, which propel the cell forward (reviewed in (Paluch et al., 2016)). In most mammalian cells, Myosin is mostly present in two isoforms: Myosin IIA and IIB (Vicente-Manzanares et al., 2009). Both, Myosin IIA and IIB, have been shown to be involved in mesenchymal migration, with Myosin IIA affecting mainly protrusion dynamics, and Myosin IIB controlling cell polarisation (Vicente-Manzanares et al., 2008) and migration speed (Betapudi et al., 2006, Halder et al., 2019). Additionally, recent studies showed that Myosin IIA generates cortical tension and controls bleb retraction during cell division (Taneja and Burnette, 2019, Taneja et al., 2020). However, the specific functions of Myosin IIA and IIB during amoeboid migration have, to our knowledge, not yet been investigated.

While mouse ES cells have been reported to display some level of motility *in vitro* (Strawbridge et al., 2020, Turco et al., 2012), their migration modes and the underlying mechanisms, and how these might change as ES cells start differentiating, have not been investigated. Here, we investigate whether the shape change characterizing early differentiation in mouse ES cells is associated with a change in the cells’ motile potential. We first quantitatively characterize the shape changes displayed by ES cells during exit of the ES cell state. We then show that naïve ES cells display limited movement on 2D substrates, but can effectively migrate in an amoeboid fashion using bleb-like protrusions when placed in confinement. We further show that cells exiting the ES cell state progressively acquire the potential to efficiently migrate on a 2D surface, using a mesenchymal-like migration mode. Interestingly, we find that confined migration of naïve ES cells strongly depends on Myosin IIA but not Myosin IIB and is enhanced by inhibition of the Arp2/3 complex. In contrast, the mesenchymal-like migration of cells exiting the ES cell state strongly depends on Arp2/3 activity and is enhanced upon Myosin IIA depletion. Together, our findings show that early differentiation of mouse ES cells is associated with extensive changes in cell migration potential, potentially caused by a switch in the regulation of the actomyosin cytoskeleton.

## Results

### Exit from ES cell state is associated with cell spreading

To investigate the changes in cell behaviour associated with exit from ES cell state, we first quantitatively characterized the morphological changes displayed during the process. ES cells were cultured in N2B27 medium supplemented with Leukaemia inhibitory factor (Lif) and two inhibitors, the Mek inhibitor PD0325901 and the Gsk3 inhibitor Chiron (2i+Lif culture medium). Exit from ES cell state was triggered by placing cells in N2B27 medium alone. Cells after 24h in N2B27 (hereafter T24 cells) have been shown to be in a transition state, where they are exiting the ES cell state, and after 48h (T48 cells), the majority of the cells have successfully exited the ES cell state and are irreversibly committed to differentiation (Kalkan et al., 2017) (Fig. 1A). For cells cultured on gelatine-coated substrates, cell spreading has been reported to occur in an asynchronous manner starting from around ~12h after triggering ES cell state exit (De Belly et al., 2020), but to what extent the shapes of individual cells differ at different stages of early differentiation has not been investigated. To address this, we quantified the shapes of ES, T24 and T48 cells plated on gelatine and expressing the membrane marker Gap43-mCherry (Fig. 1B). To facilitate shape analysis, we focused on individual cells with no neighbours, and segmented the cell’s largest crosssection (the plane at the interface with the substrate). We then compared general shape features between the different cell states (Fig. 1C, D; Fig. S1A). We analysed the cells’ cross-sectional area, as a first-degree readout of cell spreading, and solidity, which is a measure of the roughness of the cell perimeter (a solidity of 1 describes a fully convex shape, whereas values below 1 correspond to more irregularly shaped cells, with concave regions that would for example result from membrane protrusions). We found that the mean cross-sectional area increased quickly upon exit from ES cell state, (Fig. 1C; Fig. S1A). Concomitantly, solidity progressively decreased from ES cells to T24 cells an dT48 cells (Fig. 1D; Fig. S1A), indicating that spreading was associated with increased protrusive activity during exit from ES cell state.

**Figure 1:**
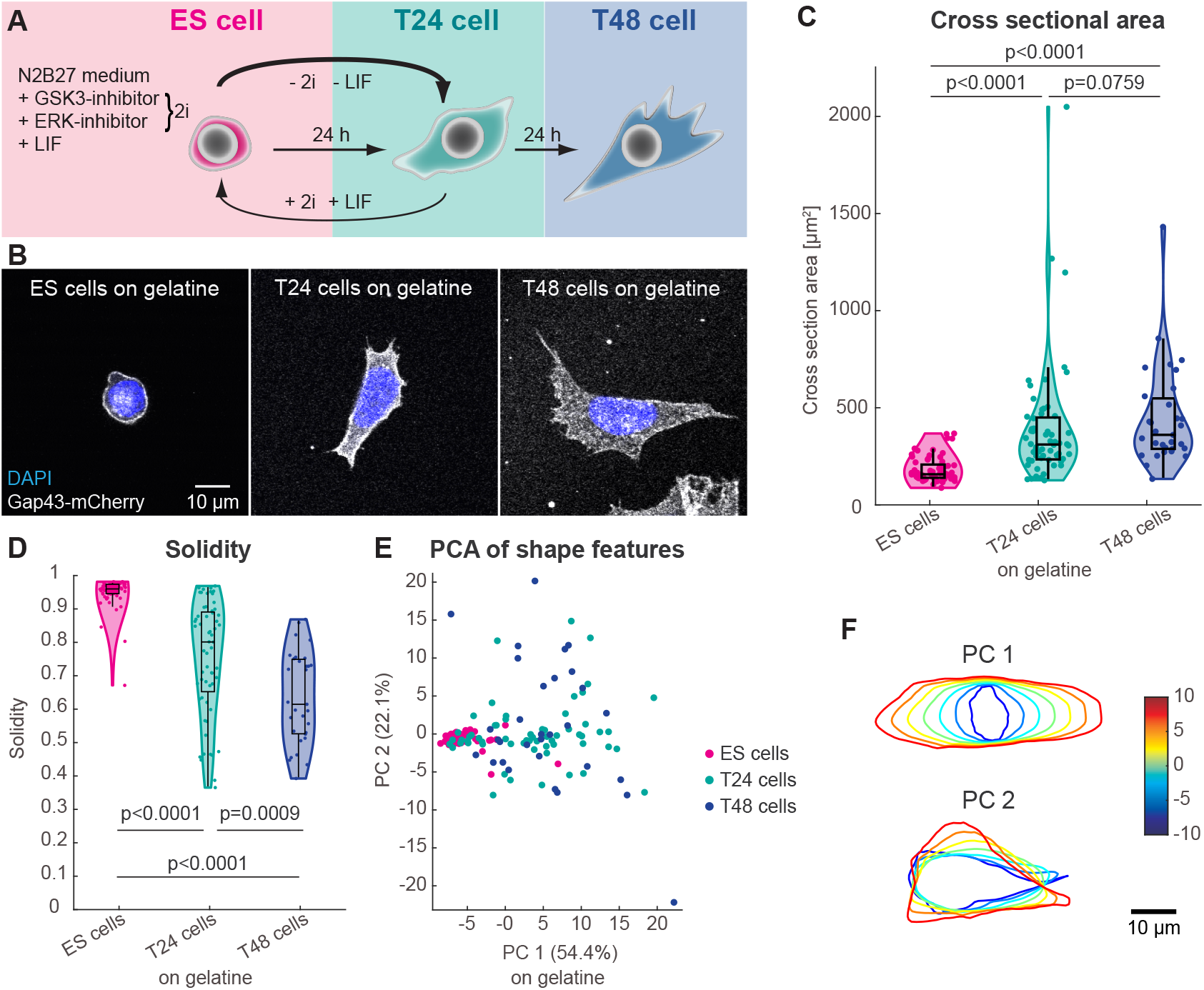
**(A)** Schematic of ES cell state exit *in vitro*. Naive ES cells (pink) are cultured in N2B27 medium supplemented with Gsk3 and Mek inhibitors and Lif. ES cells display a round morphology. When removing the two inhibitors and Lif cells exit the ES cell state and change their shape after 24h (T24 cells, green), a partially reversible process if both inhibitors and Lif are added back. After further 24h T48 cells (blue) are irreversibly committed and display a more mesenchymal-like shape. **(B)** Representative confocal images of a Rex1-GFP Gap43-mCherry cells as naïve ES cells, T24 cells and T48 cells on gelatine. One imaging plane at the interface with the substrate is shown. The nucleus is dyed using DAPI (blue), Gap-43 marks the cell membrane (grey). **(C-F)** Shape measurements of Rex1-GFP and Gap43-mCherry expressing naïve ES cells (pink; n=65, N=3), T24 cells (green; n=60, N=3) and T48 cells (blue; n=31, N=3) on gelatine including all data points. Showing **(C)** the median cross-sectional area: 157 ± 64 μm^2^ (n=65) for ES cells to 311 ± 306 μm^2^ (n=60) for T24 cells and 360 ± 256 μm^2^ (n=31) for T48 cells **(D)** Median solidity: 0.962 ± 0.046 (n=65) for ES cells, to 0.803 ± 0.170 (n=60) for T24 cells and 0.617 ± 0.136 (n=31) for T48 cells, **(E)** Principal Component Analysis (PCA) of the Fourier descriptors and **(F)** shape changes due to different values of the first and second principal component. The reference shape is the average of all cells. The colour code marks the value of the principal components, PC1 and PC2. Statistical significance in **(C** and **D)** was assessed using a Mann-Whitney U-test.

To analyse the cell shape associated with early ES cell differentiation in a more unbiased and systematic manner, we quantified segmented cell shapes using a series of harmonic functions called Fourier descriptors (Diaz et al., 1989, Lestrel, 1997). In this analysis, cell shape is described through a high-dimensional set of features, which correspond to coordinates in the Fourier descriptor space (see Methods). To represent the resulting high-dimensional data set in a low-dimensional space, we then performed a principal component analysis (PCA, see Methods). The PCA showed that 76.5% of the variability in cell shape features could be described with the two first PCs (Fig. S1B). In the corresponding 2D projection, the three cell populations (ES, T24 and T48 cells) formed two distinct clusters, with most of the ES cells in one cluster and most of the T48 cells in the other; T24 cells represented an intermediate population, with a majority of T24 cells displaying shapes similar to T48 cells, but a small sub-population displaying shapes similar to ES cells (Fig. 1E). The primary axis of the ordering of the clusters was along the direction of the first principal component (PC 1), which corresponds to an increase of area and elongation for increasing values of PC1 (Fig. 1F). ES cells had small and round cross-sectional areas, whereas T24 and T48 displayed larger areas and more elongated shapes (Fig. 1E,F). The second dominating PC, PC2, can be interpreted as describing the presence of protrusion-like elongations for increasing absolute values (Fig. 1F). While ES cells displayed values around 0 for PC2, corresponding to a lack of cellular protrusions, T24 and T48 showed a broad distribution of PC2 values (Fig. 1E,F). Together, our unbiased analysis shows that the shapes of ES, T24 and T48 cells can be described by two populations differing mostly by the cross-sectional area and levels of protrusivity of the cells.

Next, we asked whether clear shape changes could also be observed for cells exiting the ES cell state on laminin-coated substrates, where ES cells show stronger adhesion compared to gelatine (Tamm et al., 2013, Domogatskaya et al., 2008). Laminin is the first extra cellular matrix component detected in the early embryo (Cooper and MacQueen, 1983) and is commonly used for ES cell cultures (Boroviak et al., 2014). On laminin ES cells already display some level of spreading (De Belly et al., 2020) (Fig. S2A). Cells exiting the ES cell state on laminin displayed a clear increase in cross-sectional area in the first 24 h of exit (Fig. S2B, F) and a slight decrease in solidity, indicative of increased protrusions (Fig. S2C, F). PCA analysis of the Fourier descriptors coefficients associated with the cell shapes analysed showed that 67.5% of the variability in cell shape features could be described with the two first PCs (Fig. S1C). The PCA analysis indicated that the three cell populations overlapped much more and individual regions were less pronounced than for cells plated on gelatine (Fig. S2D, compare to Fig. 1E). Nonetheless, regions containing ES cells and T48 cells only partially overlapped (Fig. S2D) and the dominating PCs for cells on laminin qualitatively resembled those describing cell shapes on gelatine (Fig. 1F and S2E). Together, this analysis indicates that while ES cells grown on laminin are more spread out and less round than ES cells grown on gelatine, probably because of the increased adhesion to laminin, they nonetheless undergo enhanced spreading during ES cell state exit.

### Embryonic stem cells migrate in confined environments

We next investigated whether early ES cell differentiation is associated with changes in migratory behaviour. We first tested whether ES cells are able to migrate (Fig. 2). We used ES cells expressing the nuclear marker Histone H2B-mCherry (Cannon et al., 2015, Chaigne et al., 2020), and tracked the cell nuclei (Fig. 2B,C). We observed that on 2D, gelatine-coated substrates, unconfined ES cells did not migrate efficiently (Fig. 2C-F, Movie 1). However, amoeboid migration often manifests in confinement (Paluch et al., 2016, Liu et al., 2015). We thus used a microfabricated device to investigate ES cell migration under confinement. We chose two-dimensional (2D) confinement devices (Le Berre et al., 2014, Liu et al., 2015), which allow cells to migrate in the x- and y-direction but confines them under a PDMS roof of precise height in the z-direction (Fig. 2A). We compared ES cell behaviour under roof heights of 10 μm, which only slightly confines the cells, and 5 μm, which leads to more significant confinement (Fig. 2C). We observed that confinement induced efficient migration in ES cells, with bleb-like protrusions consistent with amoeboid migration, and frequent direction changes (Fig. 2B,C, Movie 2). We analysed the mean instantaneous cell velocity, as a direct readout of motility (Fig. 2D), and the distribution of the instantaneous velocities (Fig. 2E) which indicates what portion of a trajectory the cells spend actively migrating. We also computed the mean squared displacement (MSD) as a function of time interval Δt for each observed cell (Fig. 2F). The MSD is the squared net distance a cell has moved within a given time interval and gives information on how fast and persistent a cell moves. We found that confinement increased the instantaneous velocity from 0.48 ± 0.45 μm/min (mean ± standard deviation (SD), n = 51) for ES cells confined to 10 μm, to 1.16 ± 0.60 μm/min (mean ± SD, n = 18) for ES cells under 5 μm confinement (compared to 0.20 ± 0.07 μm/min, mean ± SD, n = 28, for unconfined ES cells, Fig. 2D, Table 1). In populations of cells confined to a 10 μm height, many cells still displayed very small (<0.5 μm/min) velocities (Fig. 2C,D), however this population was lost as cells moved at higher velocities under strong confinement (5 μm confinement, Fig. 2C,D). Consistently, the MSD of cells confined to a 5 μm height was considerably higher than for cells confined to 10 μm (Fig. 2F), indicating that cells migrated faster and with a higher degree of directionality.

**Table 1:** List of all mean instantaneous velocities and the corresponding standard deviation, including the number of cells analysed (n) and the number of biological replicates performed (N).

| Cell line | Cell state | Conf. | Drug treatment | Coating | n | N | Mean Inst. Velo. + SD |
| --- | --- | --- | --- | --- | --- | --- | --- |
| H2B-mCherry | ES cells | - | - | gelatine | 28 | 2 | 0.20 ± 0.07 µm/min |
| H2B-mCherry drift corr. | ES cells | - | - | gelatine | 28 | 2 | 0.20 ± 0.06 µm/min |
| H2B-mCherry | ES cells | - | - | laminin | 129 | 2 | 0.37 ± 0.23 µm/min |
| H2B-mCherry | ES cells | 10 µm | - | BSA | 51 | 2 | 0.48 ± 0.45 µm/min |
| H2B-mCherry drift corr. | ES cells | 10 µm | - | BSA | 51 | 2 | 0.48 ± 0.44 µm/min |
| H2B-mCherry | ES cells | 5 µm | - | BSA | 18 | 2 | 1.16 ± 0.60 µm/min |
| H2B-mCherry drift corr. | ES cells | 5 µm | - | BSA | 18 | 2 | 1.12 ± 0.58 µm/min |
| H2B-mCherry 1h N2B27 | ES cells | 5 µm | - | BSA | 42 | 2 | 0.97 ± 0.47 µm/min |
| H2B-mCherry | ES cells | 5 µm | DMSO | BSA | 22 | 2 | 0.87 ± 0.66 µm/min |
| H2B-mCherry | ES cells | 5 µm | 5 µM Blebbistatin | BSA | 25 | 2 | 0.25 ± 0.39 µm/min |
| H2B-mCherry | ES cells | 5 µm | 5 µM Y27632 | BSA | 39 | 2 | 0.15 ± 0.08 µm/min |
| H2B-mCherry | ES cells | 5 µm | 100 µM CK666 | BSA | 26 | 2 | 1.16 ± 0.87 µm/min |
| H2B-mCherry | T24 cells | 5 µm | - | BSA | 60 | 2 | 0.52 ± 0.36 µm/min |
| Myosin control | ES cells | 5 µm | - | BSA | 25 | 2 | 1.68 ± 1.46 µm/min |
| Mh9 KO | ES cells | 5 µm | - | BSA | 32 | 2 | 0.28 ± 0.15 µm/min |
| Mh10 KO | ES cells | 5 µm | - | BSA | 20 | 2 | 1.31 ± 0.94 µm/min |
| H2B-mCherry | T24 cells | - | - | gelatine | 23 | 3 | 0.69 ± 0.29 µm/min |
| H2B-mCherry | T24 cells | - | DMSO | gelatine | 256 | 8 | 0.70 ± 0.30 µm/min |
| H2B-mCherry | T24 cells | - | 5 µM Blebbistatin | gelatine | 204 | 3 | 0.90 ± 0.40 µm/min |
| H2B-mCherry | T24 cells | - | 5 µM Y27632 | gelatine | 82 | 4 | 0.80 ± 0.36 µm/min |
| H2B-mCherry | T24 cells | - | 100 µM CK666 | gelatine | 118 | 5 | 0.50 ± 0.20 µm/min |
| Myosin control | T24 cells | - | - | gelatine | 13 | 3 | 0.42 ± 0.28 µm/min |
| Mh9 KO | T24 cells | - | - | gelatine | 15 | 3 | 0.77 ± 0.40 µm/min |
| Mh10 KO | T24 cells | - | - | gelatine | 9 | 3 | 0.52 ± 0.19 µm/min |
| H2B-mCherry | T24 cells | - | - | laminin | 136 | 3 | 0.68 ± 0.38 µm/min |
| H2B-mCherry | T24 cells | - | DMSO | laminin | 148 | 8 | 0.73 ± 0.40 µm/min |
| H2B-mCherry | T24 cells | - | 5 µM Blebbistatin | laminin | 160 | 3 | 1.00 ± 0.40 µm/min |
| H2B-mCherry | T24 cells | - | 5 µM Y27632 | laminin | 53 | 3 | 0.78 ± 0.43 µm/min |
| H2B-mCherry | T24 cells | - | 100 µM CK666 | laminin | 60 | 5 | 0.46 ± 0.26 µm/min |

**Figure 2:**
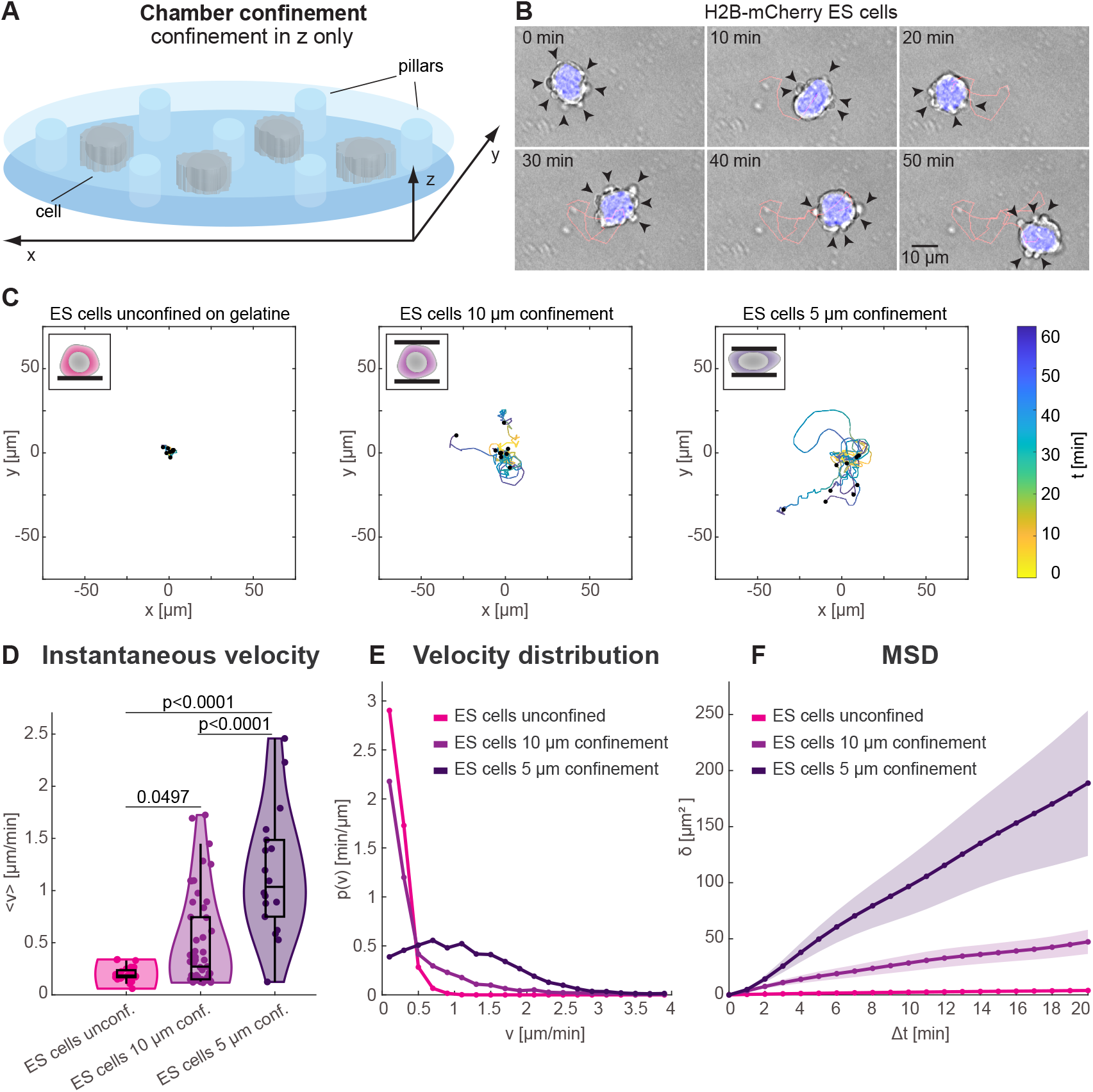
**(A)** Schematic of cell confinement between two plates kept apart by PDMS pillars. Cells are free to move in x and y directions but are confined in the z-direction. **(B)** A time course (10min interval) of a H2B-mRFP ES cell moving in 5 μm confinement. A bright field image of one plane is shown (grey), the nucleus is highlighted in blue and the cell trajectory is shown in red. Black arrows mark blebs. **(C)** Trajectories of unconfined ES cells on gelatine coating, ES cells confined under a 10 μm roof and ES cells confined under a 5 μm roof. The first 60 minutes of each trajectory are shown and 9 representative trajectories for each condition. The colour gradient is dependent on time [t]. **(D)** Violin plot of the mean instantaneous velocity of unconfined ES cells on gelatine coating (pink; 0.20 ± 0.07 μm/min; n=28, N=2)), ES cells confined under a 10 μm roof (purple; 0.48 ± 0.45 μm/min; n=51, N=2) and ES cells confined under a 5 μm roof (violet; 1.16 ± 0.60 μm/min; n=18, N=2) including all data points. Statistical significance was assessed using a Mann-Whitney U-test. **(E)** Instantaneous velocity distribution of ES cells on gelatine coating (pink), ES cells confined under a 10 μm roof (purple) and ES cells confined under a 5 μm roof (violet). **(F)** Mean squared displacement for ES cells on gelatine coated plates (pink), ES cells confined under a 10 μm roof (purple) and ES cells confined under a 5 μm roof (violet) including the mean standard error.

To confirm that our results were not affected by hydrodynamic flows in the microfabricated devices, we estimated and subtracted the linear drift of each trajectory (see methods) (Fig. S3A-C). We found that all three migration parameters analysed were not affected by flows acting on the cells (Fig. S3A-C, compare to Fig. 2D-F, Table 1). We then asked whether the lack of effective migration on 2D substrates was simply due to the cells’ round shapes when cultured on gelatine. We thus assessed cell migration on laminin, where ES cells display limited spreading (Fig. S2). Interestingly, we found that ES cells on laminin displayed a slightly more migratory behaviour on 2D substrates (instantaneous velocity 0.37 ± 0.23 μm/min, mean ± SD, n = 129) than their counterparts on gelatine, but their motility was still significantly smaller than for ES cells under confinement (Fig. S3D-G, Table 1). This suggests that spreading alone does not confer migratory potential to ES cells on a 2D substrate. Finally, we verified that the confined migration of ES cells was not affected by the presence of Lif, or Gsk-3 and Mek inhibitors in the 2i+Lif culture medium, as these inhibitors can affect the cytoskeleton (Miller et al., 2014, Sun et al., 2009). We thus placed ES cells in inhibitor-free N2B27 medium 1 h prior to the experiment and assessed their migration in confinement. We found that removing the inhibitors from the 2i+Lif culture media did not affect the migratory behaviour of cells under 5 μm confinement (Fig. S3H-K, Table 1), indicating that possible effects of the inhibitors on cytoskeletal dynamics do not significantly affect ES cell migration in confinement. Together, our observations show that ES cells are unable to migrate on 2D substrates, but display effective amoeboid-like migration when placed in confinement.

### Cells exiting the ES cell state display reduced migration in confinement, but can migrate on 2D substrates

Next, we asked if the migration potential of ES cells changes as they start to differentiate. We hypothesized that the cell spreading associated with exit of ES cell state (Fig. 1) could lead to enhanced migration on 2D substrates. We thus imaged T24 cells on 2D substrates and analysed their migratory behaviour (Fig. 3). The majority of T24 cells are found in dense colonies where individual cell migration cannot be assessed, however a subset of cells spontaneously separated from colonies (Movie 3). We tracked these transiently isolated cells and observed that they formed lamellipodia (Fig. 3A) and displayed effective migration both on gelatine- and laminin-coated substrates (Movies 3 and 4, Fig. 3B and S4A). Under these conditions, T24 cells displayed migration velocities of 0.69 ± 0.29 μm/min (mean ± SD, n = 23) on gelatine and 0.68 ± 0.38 μm/min (mean ± SD, n = 136) on laminin, thus slower than ES cells under 5 μm confinement on gelatine (Fig. 3C and S4B, Table 1). The velocity distribution and MSD also suggested that the unconfined T24 cells move slower than confined ES cells (Fig. 3D,E). Nonetheless, our results indicate that, in contrast to naïve ES cells, T24 cells efficiently migrate on 2D substrates using lamellipodia-like protrusions.

**Figure 3:**
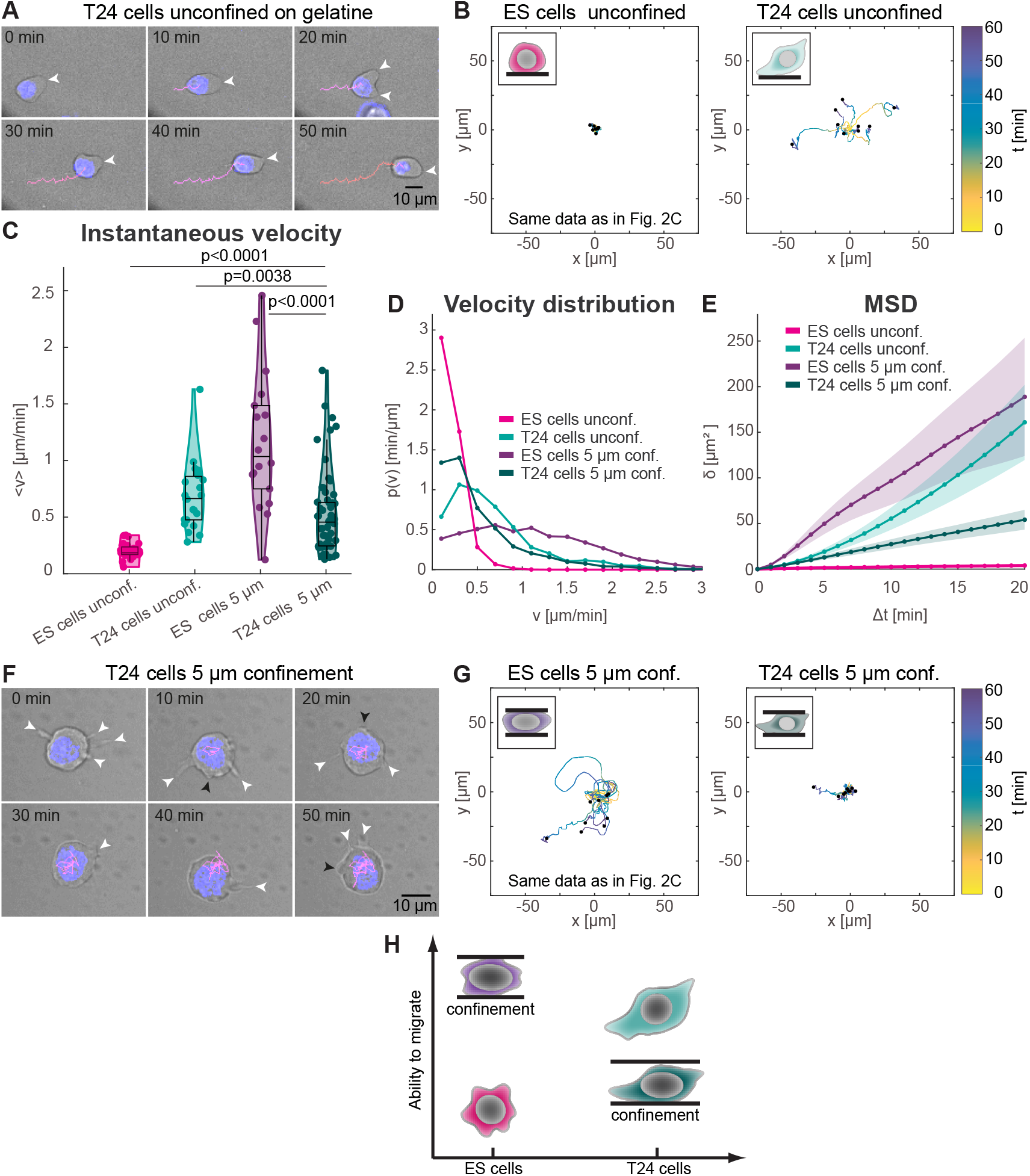
**(A)** A time course (10min interval) of an unconfined H2B-mRFP T24 cell moving on gelatine coating. A bright field image of one plane is shown (grey), the nucleus is highlighted in blue and the cell trajectory is shown in red. White arrows mark lamellipodia. **(B)** Trajectories of unconfined ES cells on gelatine coating (same data as shown in Fig. 2C) and unconfined T24 cells on gelatine coating. The first 60 minutes of each trajectory are shown and 9 representative trajectories for each condition. The colour gradient is dependent on time [t]. **(C)** Violin plot of the mean instantaneous velocity of unconfined ES cells on gelatine coating (pink; 0.20 ± 0.07 μm/min; n=28, N=2), unconfined T24 cells on gelatine coating (turquoise; 0.69 ± 0.29 μm/min; n=23, N=3), ES cells confined under a 5 μm roof (purple; 1.16 ± 0.60 μm/min; n=18, N=2) and T24 cells confined under a 5 μm roof (green; 0.52 ± 0.36 μm/min; n=60, N=2) including all data points. Statistical significance was assessed using a Mann-Whitney U-test. **(D)** Instantaneous velocity distribution of unconfined ES cells on gelatine coating (pink), unconfined T24 cells on gelatine coating (turquoise), ES cells confined under a 5 μm roof (purple) and T24 cells confined under a 5 μm roof (green). **(E)** Mean squared displacement for unconfined ES cells on gelatine coating (pink), unconfined T24 cells on gelatine coating (turquoise), ES cells confined under a 5 μm roof (purple) and T24 cells confined under a 5 μm roof (green) including the mean standard error. **(F)** A time course (10min interval) of a H2B-mRFP T24 cell in 5 μm confinement. A bright field image of one plane is shown (grey), the nucleus is highlighted in blue and the cell trajectory is shown in red. Black arrows mark blebs, white arrows mark lamellipodia. **(G)** Trajectories of ES cells confined under a 5 μm roof (same data as shown in Fig. 2C) and T24 cells confined under a 5 μm roof. The first 60 minutes of each trajectory are shown and 9 representative trajectories for each condition. The colour gradient is dependent on time [t]. **(H)** A cartoon highlighting the main findings of this figure. ES cells (pink) are unable to migrate if not confined, but start migrating in a bleb-based manner once in confinement (purple). T24 cells (turquoise) are able to migrate without confinement, but struggle to migrate under confinement (green).

We then asked if T24 cells retain the ability to migrate in confinement. To test this, we induced ES cell state exit on gelatine and after 24h confined the cells under a 5 μm roof height (Movie 5). We found that T24 cells under 5 μm confinement displayed rounded shapes with lamellipodia-like structures but also occasional blebbing (Movie 5, Fig. 3F), suggesting that T24 cells can partially switch to amoeboid-like shapes when confined. Confined T24 cells displayed a limited level of migration compared to similarly confined ES cells (Fig. 3G) or to unconfined T24 cells (Fig. 3B). Their mean instantaneous velocity was 0.52 ± 0.36 μm/min (mean ± SD, n = 60), smaller than for unconfined T24 cells and considerably smaller than for ES cells in similar confinement (Fig. 3C, Table 1). Analysis of the velocity histogram and MSD further suggested that while T24 cells maintain some migratory ability under confinement, their migration was reduced compared to similarly confined ES cells (Fig. 3C-E). Together, our data show that T24 cells retain some ability to migrate in confinement with amoeboid-like morphologies, but that cell spreading during exit from ES cell state is accompanied by acquisition of a potential for mesenchymal-like migration on 2D substrates (Fig. 3H).

### Migration of naïve ES cells in confinement strongly depends on Myosin IIA, but not on the Arp2/3 complex

Next, we sought to understand whether distinct molecular players were required for the different migration modes displayed by ES cells and cells exiting the ES cell state. We first investigated the migration of ES cells in confinement. Previous studies have shown that high Myosin II activity is a key factor promoting amoeboid migration (Liu et al., 2015) (reviewed in (Sanz-Moreno and Marshall, 2010, Agarwal and Zaidel-Bar, 2019)). To test if ES cells in confinement indeed are dependent on Myosin II, we treated cells with 5 μM of Blebbistatin, a Myosin II ATPase inhibitor (Limouze et al., 2004). We found that Blebbistatin-treated ES cells showed a clear reduction in migration velocity (0.25 ± 0.39 μm/min, mean ± SD, n = 25) compared to DMSO controls (0.87 ± 0.66 μm/min, mean ± SD, n = 22), and displayed shorter trajectories, velocity distributions shifted towards low velocities and significantly reduced MSD (Fig. 4A-D, Table 1). To further assess the dependency of ES cell migration on Myosin II, we treated cells with 5 μM of Y27632, an inhibitor of Rho-associated protein kinase (ROCK), a key activator of Myosin II (Uehata et al., 1997). ROCK inhibition almost completely suppressed ES cell migration in confinement (mean velocity: 0.15 ± 0.08 μm/min, mean ± SD, n = 39) (Fig. 4A-D, Table 1). We then asked whether confined ES cell migration would be affected by inhibition of the Arp2/3 complex, a key factor promoting mesenchymal migration and typically dispensable for amoeboid migration (Beckham et al., 2014). We thus treated ES cells with 100 μM CK666, an inhibitor of the Arp2/3 complex (Nolen et al., 2009). We observed that Arp2/3 inhibition did not inhibit the confined migration of ES cells and even led to a slight increase in cell velocities (1.16 ± 0.87 μm/min, mean ± SD, n = 26), velocity distribution and MSD (Fig. 4A-D, Table 1). Together, these data indicate that the migration of ES cells in confinement depends on Myosin II activity, but not on the Arp2/3 complex, consistent with typical features of amoeboid migration.

**Figure 4:**
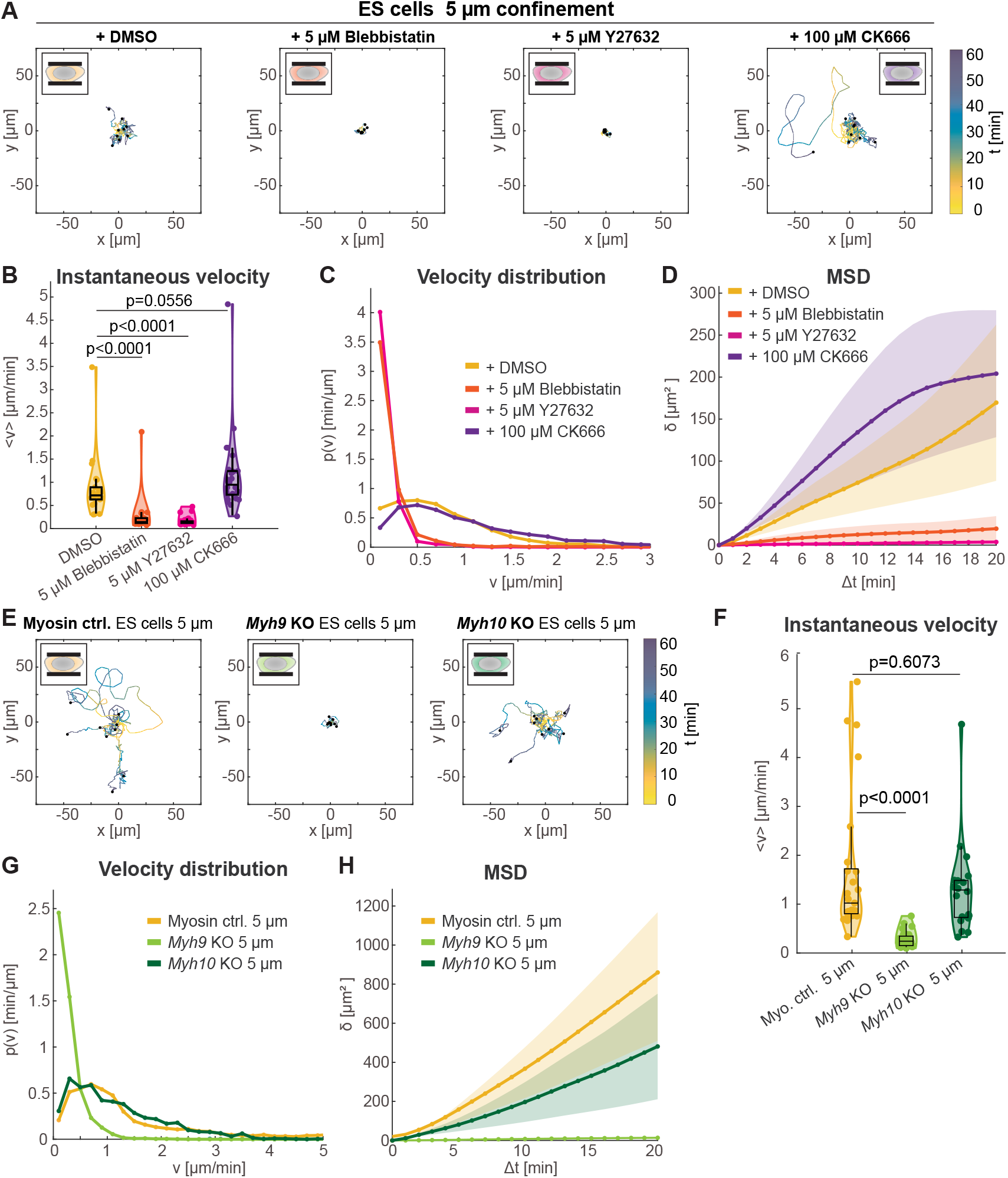
**(A**) Trajectories of 5μm confined ES cells treated with DMSO, 5 μM Blebbistatin, 5 μM Y27632 and 100 μM CK666. The first 60 minutes of each trajectory are shown and 9 representative trajectories for each condition. The colour gradient is dependent on time [t]. **(B)** Violin plot of the mean instantaneous velocity of 5μm confined ES cells treated with DMSO (yellow; 0.87 ± 0.66 μm/min; n=22, N=2), treated with 5 μM Blebbistatin (orange; 0.25 ± 0.39 μm/min; n=25, N=2), 5 μM Y27632 (pink; 0.15 ± 0.08 μm/min; n=39, N=2) and 100 μM CK666 (purple; 1.16 ± 0.87 μm/min; n=26, N=2) including all data points. Statistical significance was assessed using a Mann-Whitney U-test. **(C)** Instantaneous velocity distribution of 5μm confined ES cells treated with DMSO (yellow), 5 μM Blebbistatin (orange), 5 μM Y27632 (pink) and 100 μM CK666 (purple). **(D)** Mean squared displacement for 5μm confined ES cells treated with DMSO (yellow), 5 μM Blebbistatin (orange), 5 μM Y27632 (pink) and 100 μM CK666 (purple) including the mean standard error. **(E)** Trajectories of 5μm confined Myosin control ES cells, 5μm confined *Mhy9* KO ES cells and 5μm confined *Myh10* KO ES cells. The first 60 minutes of each trajectory are shown and 9 representative trajectories for each condition. The colour gradient is dependent on time [t]. **(F)** Violin plot of the mean instantaneous velocity of 5 μm confined Myosin control ES cells (yellow; 1.68 ± 1.46 μm/min; n=25, N=2), 5 μm confined *Mhy9* KO ES cells (light green; 0.28 ± 0.15 μm/min; n=32, N=2) and 5μm confined *Myh10* KO ES cells (dark green; 1.31 ± 0.94 μm/min; n=20, N=2) including all data points. Statistical significance was assessed using a Mann-Whitney U-test. **(G)** Velocity distribution of 5 μm confined Myosin control ES cells (yellow), 5μm confined *Mhy9* KO ES cells (light green) and 5μm confined *Myh10* KO ES cells (dark green). **(H)** Mean squared displacement for 5 μm confined Myosin control ES cells (yellow), 5μm confined *Mhy9* KO ES cells (light green) and 5μm confined *Myh10* KO ES cells (dark green) including the mean standard error.

We then asked which isoform of Myosin II drives the migration of ES cells in confinement. To address this question, we used ES cells knocked-out for either *Myh9*, the gene coding for the heavy chain of Myosin IIA, or *Myh10*, the gene coding for the heavy chain of Myosin IIB (Ma et al., 2020). We observed that when placed under 5 μm confinement, *Myh9*-deficient ES cells showed almost no motility, with strongly reduced instantaneous velocity (mean: 0.28 ± 0.15 μm/min, mean ± SD, n = 32, in *Myh9-KO* cells, compared to 1.68 ± 1.46 μm/min, mean ± SD, n = 25, in control cells (Fig. 4E-H, Table 1). In contrast, the cells’ instantaneous velocity, velocity distribution and MSD were barely affected by deletion of *Myh10* (mean instantaneous velocity of 1.31 ± 0.94 μm/min, mean ± SD, n = 20; Fig. 4E-H, Table 1).

Taken together, our data suggests that ES cells migrate in confinement using an amoeboid migration mode and show that their migration depends on Myosin IIA but not Myosin IIB.

### Migration of cells exiting the ES cell state depends on Arp2/3 activity, but not on Myosin II

Next, we investigated the molecular players required for the mesenchymal-like migration we observe in cells exiting the ES cell state. The Arp2/3 complex has been described as the main actin nucleator responsible for promoting the actin-network in the lamellipodium (Bailly et al., 1999). To test if T24 cell migration on 2D substrates depends on Arp2/3 activity, we treated cells with 100 μM CK666. This resulted in a significant reduction in mean instantaneous velocity (0.50 ± 0.20 μm/min, mean ± SD, n = 118) compared to DMSO controls (0.70 ± 0.30 μm/min, mean ± SD, n = 256); cells also displayed velocity distributions shifted towards low velocities and reduced MSD (Fig. 5B-D, Table 1). In contrast, T24 cells treated with 5 μM with Y27632 or Blebbistatin showed increased mean instantaneous velocities (0.80 ± 0.36, mean ± SD, n = 82 for Y27632-treated cells and 0.90 ± 0.40 μm/min, mean ± SD, n = 204, for Blebbistatin-treated cells), and their velocity distributions and MSDs showed slightly enhanced migration compared to control cells (Fig. 5B-D, Table 1). T24 cells grown on laminin displayed migration phenotypes very similar to cells grown on gelatine upon inhibition of either Arp2/3 complex activity or Myosin motor activity (Fig. S5A-D, Table 1).

**Figure 5:**
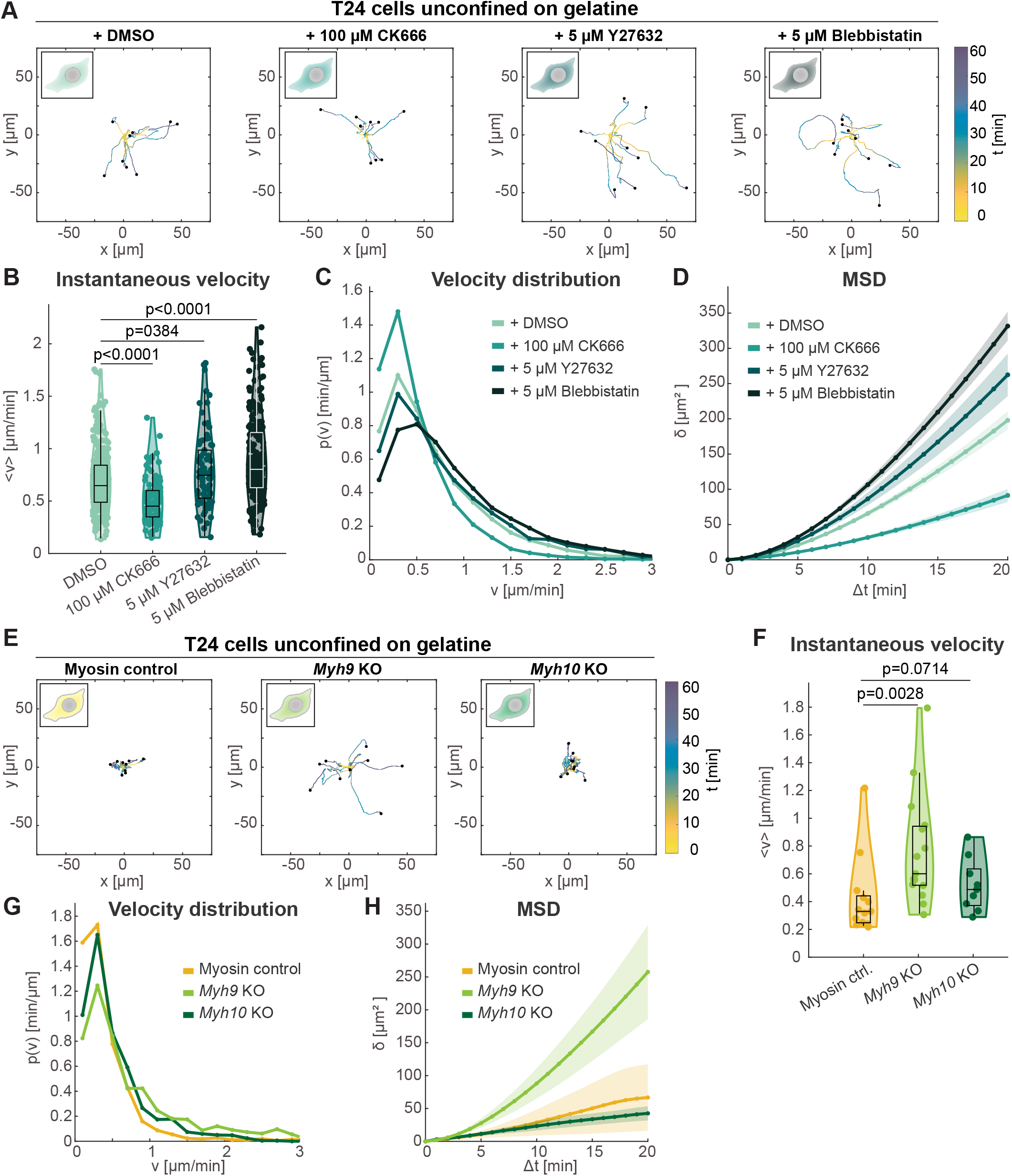
**(A**) Trajectories of unconfined T24 cells treated with DMSO, 100 μM CK666, 5 μM Y27632 and 5 μM Blebbistatin on gelatine coating. The first 60 minutes of each trajectory are shown and 9 representative trajectories for each condition. **(B)** Violin plot of the mean instantaneous velocity of unconfined T24 cells on gelatine treated with DMSO (pale green; 0.70 ± 0.30 μm/min; (n=256, N=8), treated with 100 μM CK666 (dark green; 0.50 ± 0.20 μm/min; N=118, N=5), treated with 5 μM Y27632 (grey; 0.80 ± 0.36 μm/min, (n=82, N=4) and treated with 5 μM Blebbistatin (black; 0.90 ± 0.40 μm/min; (n=204, N=3) including all data points. Statistical significance was assessed using a Mann-Whitney U-test. **(C)** Instantaneous velocity distribution of unconfined T24 cells on gelatine coating treated with DMSO (light green), 100 μM CK666 (dark green), 5 μM Y27632 (grey) and 5 μM Blebbistatin (black). **(D)** Mean squared displacement for unconfined T24 cells on gelatine treated with DMSO (light green), 100 μM CK666 (dark green), 5 μM Y27632 (grey) and 5 μM Blebbistatin (black) including the mean standard error. **(E)** Trajectories of unconfined T24 Myosin control cells, unconfined T24 *Mhy9* KO cells and unconfined T24 *Myh10* KO cells on gelatine coating. The first 60 minutes of each trajectory are shown and 9 representative trajectories for each condition. **(F)** Violin plot of the mean instantaneous velocity of unconfined T24 Myosin control cells (yellow; 0.42 ± 0.28 μm/min; n=13, N=3), unconfined T24 *Myh9* KO cells (light green; 0.77 ± 0.40 μm/min (n=15, N=3) and unconfined T24 *Mhy10* KO cells (dark green; 0.52 ± 0.19 μm/min; (n=9, N=3) on gelatine including all data points. Statistical significance was assessed using a Mann-Whitney U-test. **(G)** Velocity distribution of unconfined T24 Myosin control cells (yellow), unconfined T24 *Myh9* KO cells (light green) and unconfined T24 *Mhy10* KO cells (dark green). **(H)** Mean squared displacement for unconfined T24 Myosin control cells (yellow), unconfined T24 *Myh9* KO cells (light green) and unconfined T24 *Mhy10* KO cells (dark green) including the mean standard error.

Finally, we asked whether the slightly enhanced migration of T24 cells upon Myosin inhibition is the effect of a specific Myosin II isoform. We thus characterised migration of Myh9 and Myh10 KO cells upon exit from ES cell state. We found that T24 *Myh9* KO cells showed increased migration (0.77 ± 0.40 μm/min, mean ± SD, n = 15) compared to both Myosin control T24 cells (0.42 ± 0.28 μm/min, mean ± SD, n = 13) and T24 *Myh10* KO cells (0.52 ± 0.19 μm/min, mean ± SD, n = 9) (Fig. 5E-H, Table 1). These observations suggest that Myosin IIA, but not Myosin IIB, limits the mesenchymal-like migration of T24 cells. This suggests that Myosin IIA might be part of the molecular switch regulating the amoeboid to mesenchymal transition of ES cells during early differentiation.

Together, these results suggest that T24 cells migrate on 2D substrates using a mesenchymal-like migration mode, where migration does not strongly rely on Myosin II motors, but is enhanced by Arp/3 complex activity.

## Discussion

In this study, we investigate changes in cellular shape and migration potential as mouse ES cells exit the ES cell state. We used an automated morphometric analysis pipeline (Andrews et al., 2020, Bodor et al., 2020) to analyse cell shape in an unbiased manner. This allowed us to show that cell shape changes during exit from ES cell state can primarily be described by cell spreading, as evidenced by increased crosssectional area (Figs. S1A and S2F), and enhanced variability in cell shape due to enhances cellular protrusions (variability along the PC2 axis, Figs. 1E,F and S2D,E).

This prompted us to investigate cell migration during early differentiation, as cell spreading is a key feature of amoeboid-to-mesenchymal transitions (Pankova et al., 2010). Indeed, we found that while ES cells do not appear to be able to migrate in 2D, exit from ES cell state is associated with increased migratory potential on 2D substrates (Fig. 3A-E). Mesenchymal-like migration of cells exiting the ES cell state displayed a strong dependence on the activity of the Arp2/3 complex, consistent with the finding that Arp2/3 is essential for the mesenchymal migration of fibroblasts (Wu et al., 2012, Suraneni et al., 2012). In contrast, 2D migration of cells exiting the ES cell state did not require Myosin II activity, and migration velocity was even enhanced upon Myosin inhibition and in cells KO for Myosin IIA. This is consistent with observations in other systems, which had shown that reducing Myosin II levels or activity either does not affect or increases the speed of mesenchymal migration (Lo et al., 2004, Singh et al., 2020).

Interestingly, naïve cells were able to efficiently migrate when placed in confinement (Fig. 2). Interestingly, confined amoeboid-like migration of ES cells displayed a strong dependence on Myosin IIA but not Myosin IIB (Fig. 4). This finding is in line with the observation that Myosin IIA is necessary for the fast amoeboid motility of T-cells (Jacobelli et al., 2009), which is confinement-dependent (Jacobelli et al., 2010). Naïve ES cell migration was also slightly enhanced upon inhibition of the Arp2/3 complex. We have previously shown that Arp2/3 inhibition can lead to higher cortical tension (Bergert et al., 2012) and, as a result, would act to promote blebbing (Langridge and Kay, 2006) and amoeboid-like migration (Obeidy et al., 2020). Together our findings are consistent with previous observations in other cellular systems, that an increase in cell contractility and cortex tension favours fast amoeboid-like migration of ES cells. How exactly the transition from amoeboid-to mesenchymal-like motility is regulated during early differentiation of ES cells remains to be investigated. Sequencing data show a strong decrease in Myosin IIA transcription levels during exit from the ES cell state and an overall lower level of Myosin IIB ((Kalkan et al., 2017), Table S1); levels of Arp2/3 components however display an overall decrease during exit from ES cell state ((Kalkan et al., 2017), Table S1). As the activities of Myosin motors and the Arp2/3 complex are controlled by multiple upstream regulators (Vicente-Manzanares et al., 2009, Goley and Welch, 2006), it is likely that the machinery controlling migration during ES cell early differentiation is controlled via regulation of activity, rather than simply via expression levels of key direct effectors.

In summary, our study identifies a transition in migration mode as ES cells exit the ES cell state. It will be interesting to investigate to what extent and at what timepoints such transitions occur in development, and to further explore which upstream regulators control cell shape and migratory behaviour during early differentiation.

## Materials and methods

### ES cell lines and culturing of ES cells

Mouse ES cells were routinely cultured as described in (Mulas et al., 2019) using N2B27 2i + Lif medium at 37°C and 7% CO^2^ in culture flasks coated with 0.1% gelatine (Sigma-Aldrich, #G1890 SIGMA) diluted in PBS. N2B27 was made using DMEM/F-12 (Sigma-Aldrich, #D6421-6) and Neurobasal medium (Life Technologies, #21103-049) in a 1:1 mixture, 2.2 mM L-Glutamine (Life Technologies, #25030-024), 1% batch tested B27 (Life Technologies, #12587010), 1.33% BSA (Thermo Fisher Scientific, #15260037), 50 mM β-Mercaptoethanol (Sigma-Aldrich, #3148-25ML),12.5 ng/ml Insulin zinc (Sigma-Aldrich, #I9278) and 0.5% home-made N2. The 200x home-made N2 was made using 0.791 mg/ml Apotransferrin (Sigma-Aldrich, #T1147), 1.688 mg/ml Pu-trescine (Sigma-Aldrich, #P5780), 3 μM Sodium Selenite (Sigma-Aldrich, #S5261), 2.08 μg/ml Progesterone (Sigma-Aldrich #P8783) and 8.8% BSA (Thermo Fisher Scientific, #15260037). To culture naive ES cells in N2B27 2i + Lif 3 μM Chiron (Cambridge Bioscience, #CAY13122), 1 μM PD0325901 (Sigma-Aldrich, #PZ0162) and 10 ng/ml Lif (Merck Millipore, #ESG1107) were added to the culture medium. ES cells were passaged every other day using Accutase (Sigma-Aldrich, #A6964) and regularly tested for mycoplasma.

In this study we use E14 cells stably expressing H2B-mRFP (Cannon et al., 2015, Chaigne et al., 2020). *Myh9* KO ES cells, Myh10 KO ES cells and Myosin control ES cells from the same background (v6.5 ES cell line) were a kind gift from Robert S. Adelstein and Xuefei Ma (National Institutes of Health) (Ma et al., 2020). E14 cells stably expressing Rex1-GFP and Gap43-mCherry were a kind gift from Carlas Mulas (Stem Cell Institute, University of Cambridge).

### Drug treatment

The following inhibitors were used on ES cells and T24 cells. 5 μM Y-27632 ROCK inhibitor (Tocris, #1254), 5 μM Blebbistatin Myosin-Inhibitor (Sigma Aldrich, #B0560-1MG) and 100 μM Arp2/3 inhibitor CK-666 (MERCK Millipore, #182515-25MG) both confined and unconfined cells were incubated with the drugs for two hours before imaging was started. All drugs were also added to the imaging medium and exposure to blue light was avoided for all. DMSO was used as control, the concentration was matched to the highest concentration used in combination with a drug.

### Exit from ES cell state and imaging of cells without confinement

Exit from ES cell state was triggered by passaging the cells and seeding them in N2B27 alone. Cells were directly plated into 8 well slides (ibiTreat, #80826) in N2B27 medium. The 8 well dishes were plasma activated for 30 seconds and coated with either 0.1% gelatine (Sigma-Aldrich, #G1890 SIGMA) in PBS or 20 μg/ml laminin (Merck Millipore, #CC095) in PBS for 4h at 37°C. The cells were seeded in N2B27 medium and incubated for 24h. ES cells without confinement were plated in N2B27 2i + Lif medium the same way as T24 cells. Myosin II knockout cells and control cells were stained with 1:10,000 Cell Mask Orange (Life Technologies, #C10045) which was added 30min before imaging was started.

Imaging was performed on either an inverted confocal microscope Olympus FluoView FV1200 using a 60x silicon lens (UPLSAPO60XS) or on a widefield inverted Nikon TiE microscope using a 40x plan apo air lens. A frame rate of 1 frame/minute was applied for up to 800 minutes and the cells were kept at 37°C and 5% CO^2^ the entire time.

### Confinement and imaging of cells

ES cells for confinement experiments were grown in 0.1% gelatine (Sigma-Aldrich, #G1890 SIGMA) in PBS coated T12.5 flasks using N2B27 2i + Life medium. T24 cells for confinement experiments were exited either in a 0.1% gelatine coated T12.5 flasks or in a 20 μg/ml laminin (Merck Millipore, #CC095) coated T12.5 flask incubated for 3h at 37°C, in N2B27 medium for 24h.

We used the two-dimensional cell confiner developed in the lab of Matthieu Piel (Le Berre et al., 2014). The suction cup was made as described by (Le Berre et al., 2014) from Polydimethylsiloxane (PDMS) a silicon elastomer (Univar, #184S1.1) in a 1:10 ratio using an aluminium mould kindly milled by Andrew C. Graham. The silicon wafer used to make cover slips with a 5 μm roof height was kindly made by Ravi Desai and the Making STP (The Francis Crick Institute), the silicon wafer used to make coverslips with a 10 μm roof height was kindly lent to us by the Piel lab (Institut Curie). Cover slips bearing the PDMS pillars structures to create a precise roof height were made as described in (Le Berre et al., 2014) using PDMS in a 1:10 ratio. 50 mm FluoroDish tissue culture dishes (World Precision Instruments, Inc., #FD5040-100) were coated with 1:10 PDMS using a spin coater. The PDMS coated surface of the 50 mm FluoroDish dish as well as the side of the coverslip bearing the PDMS pillars structures were incubated with 100 μg/ml BSA (Sigma-Aldrich, #A9647-100G) in PBS for 30 min at room temperature. Subsequently both surfaces were incubated with culture medium (N2B27 alone or N2B27 2i + Lif) for at least 2h at 37°C before cells were added. Cells were detached using Accutase (Sigma-Aldrich, #A6964). For Myosin II knockout cells and control cells 1:10,000 Cell Mask Orange (Life Technologies, #C10045) was added to the detached cells. A cell suspension containing 7.5 x 10^5^ cells was added to the 50 mm FluoroDish already mounted on the microscope. The suction cup was cleaned with 70%ethanol, dried and plugged to a precise vacuum generator (Elveflow, AF1 dual pump). The non-structured side of the Coverslip was cleaned and dried before being stuck to the central column of the suction cup with the feature bearing side facing away from the suction cup. The assembled device was carefully put on the culture dish containing the cell suspension. Immediately a negative pressure of −30 mbar was applied to attach the suction cup to the dish, without confinement being applied to the cells. Subsequently the pressure was slowly raised to −90mbar, which was sufficient to pull the central column with the coverslip attached down on the culture dish, confining the cells. Imaging was performed on an inverted confocal microscope Olympus FluoView FV1200 using a 60x silicon lens (UPLSAPO60XS) and a frame rate of 1 frame/minute was applied for up to 800 minutes. Throughout the imaging process cells were kept at 37°C and 5% CO^2^.

### Image processing, segmentation and tracking

Time lapse images acquired on a widefield inverted Nikon TiE microscope were corrected for stage drift using the fluorescent channel and the Fiji (Schindelin et al., 2012) plug in StackReg (Thevenaz et al., 1998). The H2B-mRFP staining for ES cells and T24 cells, of both confined and unconfined cells, was used for tracking nuclei. For Myosin II knockout and control cells the Cell Mask Orange staining was used for tracking the cell membrane. Tracking was performed using the Fiji (Schindelin et al., 2012) plugin TrackMate (Tinevez et al., 2017). For all cells the Log detector was used with an estimated blob diameter set to 23μm for H2B-mRFP cells in confinement and without confinement. For Myosin II knockout and control cells in confinement an estimated blob diameter of 25μm was used and for non-confined cells 40μm was used. For all conditions only single cells were tracked and tacks of colonies were excluded manually. Tracks were manually stopped if a cell divided or collided with another cell. All spot statistics were exported and subsequently used for statistical analysis.

### Statistical analysis of trajectories

We estimated the instantaneous velocity of each cell by calculating the net distance that a cell has travelled within a 5 minute interval. Then, we estimated the mean instantaneous velocity for each cell by time-averaging the instantaneous absolute velocities.

The trajectories of cells were analysed with *Matlab R2019b* on the operating system *MacOS 10.15.* To account for rare time points where the cells could not be detected or tracked, the trajectories were linearly interpolated with a time interval of 1 minute. We only analysed trajectories of a total duration of at least 60 minutes. For confined migration, we only picked trajectories that end within the first 120 minutes from the start of the experiment (T0).

The instantaneous velocities of cells were computed with the equation

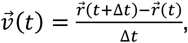

with the two-dimensional cell positions *r*(*t*) at time *t*. The mean instantaneous velocity of each trajectory was calculated by time-averaging its absolute velocities. The violin plots and boxplots in Fig. 2D, 3C, 4B, 4F, 5B, 5F, S3A, S3E, S3I, S4B and S5B were created with the help of the data visualization toolbox *Gramm* (Morel, 2018). The bandwidth of the violin plots is 0.35 μm/min. To test the significance between the mean instantaneous velocities we used a Mann-Whitney U-test.

To generate the probability distributions of the instantaneous velocity in Fig. 2E, 3D, 4C, 4G, 5C, 5G, S3B, S3F, S3J, S4C and we first created the normalized histograms of instantaneous velocity of each individual trajectory with a bin width of 0.2 μm/min. We then averaged the individual histograms for each condition.

The time-averaged mean squared displacement (MSD) of an individual trajectory, shown in Fig. 2F, 3E, 4D, 4H, 5D, 5H, S3C, S3G, S3K, S4D and S5D was computed with the equation

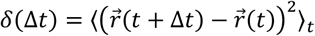

for time intervals Δ*t* ranging from 0 to 20 minutes. The MSDs for each condition were then averaged over individual trajectories. The error bars of the mean MSD correspond to the standard error of the MSD of the individual trajectories.

We also checked the impact of a linear drift due to hydrodynamic flows affecting the trajectories of the cells that might have been missed by the StackReg drift correction. To this aim, we computed the time-averaged mean displacement 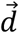

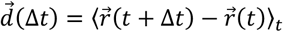

and fitted a linear function of the form 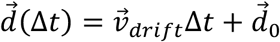, with the time-independent drift velocity 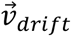, for each trajectory. We then subtracted the linear drift 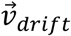 from each trajectory,

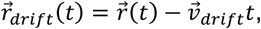

and calculated the mean instantaneous velocity, the time-averaged mean squared displacement and the probability function of instantaneous velocities from the new cell position 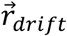.

### Shape analysis and dimensional reduction

#### Cell preparation and imaging

For shape analysis E14 cells stably expressing Rex1-GFP and Gap43-mCherry a kind gift from Carlas Mulas (Stem Cell Institute, University of Cambridge) were used. ES cells, T24 cells and T48 cells were all plated at the same time using 50% of the cell material for T48 cells in a 35mm Ibidi dish (ibiTreat, #81156) in N2B27 medium or N2B27 2i + Lif medium for ES cells. The 35mm dishes were plasma activated for 30 seconds and coated with either 0.1% gelatine (Sigma-Aldrich, #G1890 SIGMA) in PBS or 20 μg/ml laminin (Merck Millipore, #CC095) in PBS for 4h at 37°C. ES cells and T24 cells were fixed after 24h and T48 cells were fixed after 48h using 4% PFA (Fisher Scientific, #11586711) for 10 minutes at room temperature. Subsequently, cells were stained with DAPI (Sigma, #D9542-10MG) used at 10 μg/ml in PBS for 10 min at room temperature. The cells were washed three times with PBS containing 0.1% Tween-20 (Sigma-Aldrich, #P1379-500ML) for 10 min and kept in PBS until imaging was performed. Imaging was performed on a Leica TCS SP5 inverted confocal microscope using a 63x oil lens (HCX PL APO) a single image of a cell at the interface with the substrate was acquired using identical settings for each sample within the biological replicate.

#### Image processing and segmentation

The segmentation of the microscopy data of cells were performed on a *Windows 10* computer in *Fiji* (Schindelin et al., 2012), using the *Trainable Weka segmentation* plugin (Arganda-Carreras et al., 2017). For each case (laminin and gelatine) and each time point (ES, 24h and 48h) we trained a classifier on four, randomly chosen microscopy images of the cell membranes. We used a fast-random forest classifier, initialized with 100 trees and considering five different types of features: Gaussian Blur, Sobel Filter, Hessian, Difference of Gaussians and Membrane Projections. We found that the Weka segmentation is superior to normal thresholding of the fluorescence images since it accounts more reliably to membrane protrusions such as filopodia. After applying the appropriate classifiers to all experimental data, we manually check that the segmentation worked correctly and removed data where this is clearly not the case. Finally, we get a probability map that gives each pixel a probability to be within the cell membrane or nucleus.

To generate the binary shape from the probability map, we use *Matlab R2019b* on *MacOS 10.15*. To this aim, we use the built-in functions of *Matlab* and threshold the probability map with Otsu’s method and fill holes of the resulting binary shape.

### Shape quantification, statistical analysis and principal component analysis

The two-dimensional binary shapes of the cells were analysed using built-in functions of *Matlab R2019b* on *MacOS 10.15*. From the binary shapes, we compute the crosssectional area and the solidity. Solidity is a measure of the roughness of the cell perimeter by calculating the ratio of cross section area of the cell and the cross-section area of its convex hull.

The violin plots and boxplots in Fig. 1C, 1D, S1B were created with the help of the Matlab data visualization toolbox *Gramm* (Morel, 2018). To set an appropriate bandwidth of the shape feature violin plots, we computed the standard deviation (SD) of the corresponding features of all cells (independent of the time point) and set the bandwidth to half of the standard deviation of the respective data. The significance of the difference of distributions of shape features at different time points was calculated with a Mann-Whitney U-test.

Before we estimated the Fourier descriptor to the cell perimeter, we first aligned the cells. To this aim, we positioned the centre of the cell perimeter at the origin of a cartesian coordinate system and fitted an ellipse to the cell boundary. We then rotated the cell such that the long axis of the ellipse points in the x-direction of the coordinate system. We also took care that the third moment in the x-direction is positive and for cases where that was not the case, we rotated the cell 180°. Additionally, we made corrections to the perimeter coordinates by taking care that they are ordered in a clockwise manner and by choosing the point that is closest to the x-axis and has a positive x value. These corrections are based on (Sanchez-Corrales et al., 2018). With the perimeter coordinates *x_p_*(*s*) and *y_p_*(*s*), where *s* ∈ [0,1], we can then write the Fourier series up to an order *n_max_*

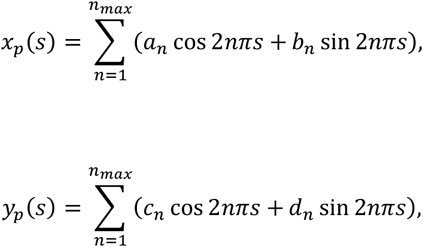

with the features *a_n_, b_n_, c_n_* and *d_n_* of order *n*. Their derivation is described in (Diaz et al., 1989). We calculate these features for all cells up to an order *n* = 40 with a self-written Matlab script and then perform a principal component analysis with the built-in Matlab function *pca()*.

To study the impact of changes of the individual principal components on the shape (see Fig. 1F and S1E) of the cells, we first compute the average shape of all cells by computing the mean values 〈*a_n_*〉, 〈*b_n_*〉, 〈*c_n_*〉 and 〈*d_n_*〉 and then add or subtract the contributions of the principal components to see how the shape changes.

## Acknowledgements

We thank Buzz Baum and Guillaume Charras for comments on the manuscript. We thank Ravi Desai and the Making STP (The Francis Crick Institute) for making silicon wafers. We thank Matthieu Piel and Juan Manuel García Arcos (Institut Curie) for intellectual input and help with the confinement device. We thank the Paluch Lab for scientific discussions and input. We thank Meghan Agnew for technical support. Furthermore, we thank the light microscopy facility at MRC-LMCB, especially Andrew Vaughan and Ki Hng for technical support. We thank Buzz Baum (MRC Laboratory of Molecular Biology) and Guillaume Charras (London Center for Nanotechnology, UCL) for input on the manuscript. We thank Carla Mulas, Celine Labouesse, and Jenny Nichols (Cambridge Stem Cell Institute) for providing reagents as well as technical expertise and advice. We thank Robert S. Adelstein and Xuefei Ma (National Institutes of Health) for sharing the Myosin knock out lines and Jonathan Chubb (MRC-LMCB) for providing the H2B-mCherry line. IMA would like to thank Trevor Graham for input on data analysis and Andrew C. Graham for milling the aluminium moulds used for making the PDMS suction cup.

## Data availability

All data and materials used and analysis in this manuscript are available upon request. All image processing and statistical analysis tools have been previously published and are publicly available.

---

**Movie 1:**

Representative bright field time lapse microscopy of a H2B-mRFP ES cell migrating on a gelatine coated 2D substrate. The nucleus is highlighted in red. Acquisition rate: 1 frame/minute. Scale bar 20 μm.

**Movie 2:**

Representative bright field time lapse microscopy of a H2B-mRFP ES cell migrating in 5μm confinement. The nucleus is highlighted in red. Acquisition rate: 1 frame/minute. Scale bar 20 μm.

**Movie 3:**

Representative bright field time lapse microscopy of a H2B-mRFP T24 cell migrating on a gelatine coated 2D substrate. The nucleus is highlighted in red. Acquisition rate: 1 frame/minute. Scale bar 20 μm.

**Movie 4:**

Representative bright field time lapse microscopy of a H2B-mRFP T24 cell migrating on a laminin coated 2D substrate. The nucleus is highlighted in red. Acquisition rate: 1 frame/minute. Scale bar 20 μm.

**Movie 5:**

Representative bright field time lapse microscopy of a H2B-mRFP T24 cell migrating on in 5 μm confinement. The nucleus is highlighted in red. Acquisition rate: 1 frame/minute. Scale bar 20 μm.

