## Supplemental Information for "Mouse embryonic stem cells switch migratory behaviour during early differentiation"

**Supplementary Figure 1:**

**(A)** Mean values for cross sectional area and solidity and the corresponding standard deviation, including the number of cells analysed (n) and the number of biological replicates performed (N). **(B, C)** Percentage of the total variance explained by each principal component (blue bars) and cumulative contribution to the total variance (red points) for the corresponding principal component analysis in **(B)** Fig. 1E and **(C)** Fig. S1D.

**Supplementary Figure 2:**

**(A)** Representative confocal images of a Rex1-GFP Gap43-mCherry cells as naïve ES cells, T24 cells and T48 cells on laminin. One imaging plane at the interface with the substrate is shown. The nucleus is dyed using DAPI (blue), Gap-43 marks the cell membrane (grey). **(B-E)** Shape measurements of Rex1-GFP and Gap43-mCherry expressing naïve ES cells (pink; n=82, N=3), T24 cells (green; n=104, N=3) and T48 cells (blue; n=43, N=3) on laminin including all data points. Showing **(B)** the median cross-sectional area: 493 ± 164 $\mu m^2$ (n=82) in ES cells to 760 ± 273 $\mu m^2$ (n=104) in T24 cells and 798 ± 301$\mu m^2$ (n=43) in T48 cells **(C)** Median solidity: 0.814 ± 0.088 (n=82) in ES cells to 0.835 ± 0.090 (n=104) in T24 and 0.791 ± 0.123 (n=43) in T48 cells, **(D)** Principal Component Analysis (PCA) of the Fourier descriptors and **(E)** shape changes due to different values of the first and second principal component. The reference shape is the average of all cells. The colour code marks the values of PC 1 and PC 2. Statistical significance in **(B** and **C)** was assessed using a Mann-Whitney U-test. **(F)** Mean values for cross sectional area and solidity and the corresponding standard deviation, including the number of cells analysed (n) and the number of biological replicates performed (N).

**Supplementary Figure 3:**

**(A)** Violin plot of the drift corrected mean instantaneous velocity of unconfined ES cells on gelatine coated plates (pink; 0.20 ± 0.06 µm/min; n=28, N=2), ES cells confined under a 10 µm roof (purple; 0.48 ± 0.44 µm/min; n=51, N=2) and ES cells confined under a 5 µm roof (violet; 1.12 ± 0.58 µm/min; n=18, N=2) including all data points. Statistical significance was assessed using a Mann-Whitney U-test. **(B)** Drift corrected instantaneous velocity distribution of ES cells on gelatine coated plates (pink), ES cells confined under a 10 µm roof (purple) and ES cells confined under a 5 µm roof (violet). **(C)** Drift corrected mean squared displacement for ES cells on gelatine coated plates (pink), ES cells confined under a 10 µm roof (purple) and ES cells confined under a 5 µm roof (violet) including the mean standard error. **(D)** Trajectories of unconfined ES cells on laminin coating, unconfined ES cells on gelatine coating (data is the same as in Fig. 2C) and confined ES cells under a 5 µm roof (data is the same as in Fig. 2C). The first 60 minutes of each trajectory are shown and 9 representative trajectories for each condition. **(E)** Violin plot of the mean instantaneous velocity of unconfined ES cells on laminin coating (green; 0.37 ± 0.23 µm/min; n=129, N=2), unconfined ES cells on gelatine coating (pink; 0.20 ± 0.07 µm/min; n=28, N=2) and ES cells confined under a 5 µm roof (violet; 1.16 ± 0.60 µm/min; n=18, N=2) including all data points. Statistical significance was assessed using a Mann-Whitney U-test. **(F)** Velocity distribution of unconfined ES cells on laminin coating (green), unconfined ES cells on gelatine coating (pink) and ES cells confined under a 5 µm roof (violet). **(G)** Mean squared displacement for unconfined ES cells on laminin coating (green), unconfined ES cells on gelatine coating (pink) and ES cells confined under a 5 µm roof (violet) including the mean standard error. **(H)** Trajectories of confined ES cells under a 5 µm roof grown in N2B27 medium alone starting 1h before imaging was started. The first 60 minutes of each trajectory are shown and 9 representative trajectories for each condition. **(I)** Violin plot of the mean instantaneous velocity of ES cells confined under a 5 µm roof height (purple; 1.16 ± 0.60 µm/min; n=18, N=2) grown in N2B27 +2i +Lif and ES cells confined under a 5 µm roof height (orange; 0.97 ± 0.47 µm/min; n=42, N=2) grown in N2B27 alone starting 1h prior imaging including all data points. Statistical significance was assessed using a Mann-Whitney U-test. **(J)** Instantaneous velocity distribution of ES cells confined under a 5 µm roof grown in N2B27 +2i +Lif (purple) and ES cells confined under a 5 µm roof grown in N2B27 (orange). **(K)** Mean squared displacement for ES cells confined under a 5 µm roof grown in N2B27 +2i +Lif (purple) and ES cells confined under a 5 µm roof grown in N2B27 (orange) including the mean standard error.

**Supplementary Figure 4:**

**(A)** Trajectories of unconfined ES cells on laminin coating (data is the same as in Fig. S3D) and unconfined T24 cells on laminin coating. The first 60 minutes of each trajectory are shown and 9 representative trajectories for each condition. **(B)** Violin plot of the mean instantaneous velocity of unconfined ES cells on laminin (purple; 0.37 ± 0.23 µm/min; n=129, N=3 and unconfined T24 cells on laminin (green; 0.68 ± 0.38 µm/min; n=136, N=3. Statistical significance was assessed using a Mann-Whitney U-test. **(C)** Instantaneous velocity distribution of unconfined ES cells on laminin (purple) and unconfined T24 cells on laminin (green). **(D)** Mean squared displacement for unconfined ES cells on laminin (purple) and unconfined T24 cells on laminin (green) including the mean standard error.

**Supplementary Figure 5:**

**(A)** Trajectories of unconfined T24 cells treated with DMSO, 100 µM CK666, 5 µM Y27632 and 5 µM Blebbistatin on laminin coating. The first 60 minutes of each trajectory are shown and 9 representative trajectories for each condition. **(B)** Violin plot of the mean instantaneous velocity of unconfined T24 cells on laminin treated with DMSO (light green; 0.73 ± 0.40 µm/min; (n=148, N=8), treated with 100 µM CK666 (dark green; 0.46 ± 0.26 µm/min; N=60, N=5), treated with 5 µM Y27632 (grey; 0.78 ± 0.43 µm/min, (n=53, N=3) and treated with 5 µM Blebbistatin (black; 1.00 ± 0.40 µm/min; (n=160, N=3) including all data points. Statistical significance was assessed using a Mann-Whitney U-test. **(C)** Instantaneous velocity distribution of unconfined T24 cells on laminin coating treated with DMSO (light green), 100 µM CK666 (dark green), 5 µM Y27632 (grey) and 5 µM Blebbistatin (black). **(D)** Mean squared displacement for unconfined T24 cells on laminin coating treated with DMSO (light green), 100 µM CK666 (dark green), 5 µM Y27632 (grey) and 5 µM Blebbistatin (black) including the mean standard error.

**Supplementary Table 1:**

**(A)** Table of ribosome-depleted (total) RNA; RPKM = reads per kilobase of transcript per million mapped reads. Data taken from (Kalkan et al., 2017).

### Supplementary Figure 1

**A**

| Cells | Cross sectional area | Solidity | n | N |
|---|---|---|---|---|
| ES | 157 ± 64 μm$^2$ | 0.962 ± 0.046 | 65 | 3 |
| T24 | 311 ± 306 μm$^2$ | 0.803 ± 0.170 | 60 | 3 |
| T48 | 360 ± 256 μm$^2$ | 0.617 ± 0.136 | 31 | 3 |

**B**

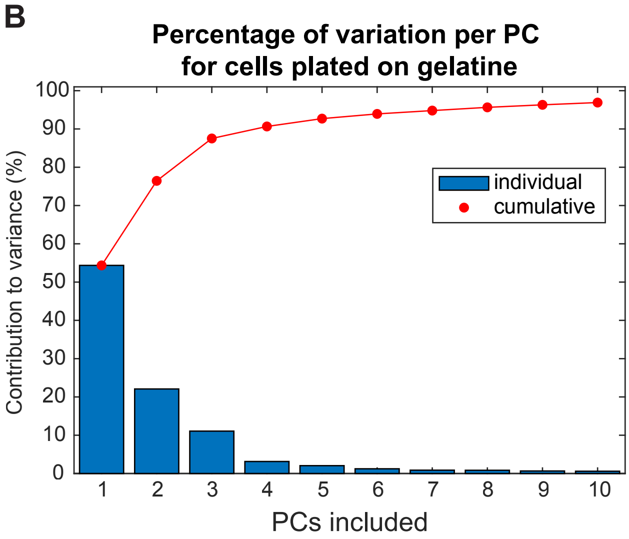

**C**

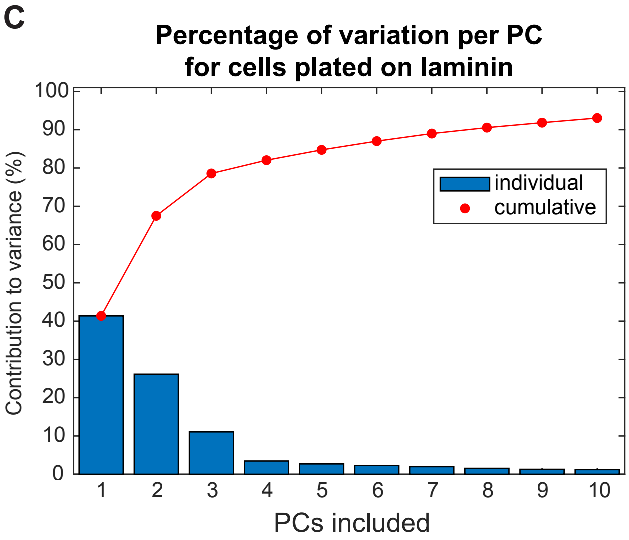

### Supplementary Figure 2

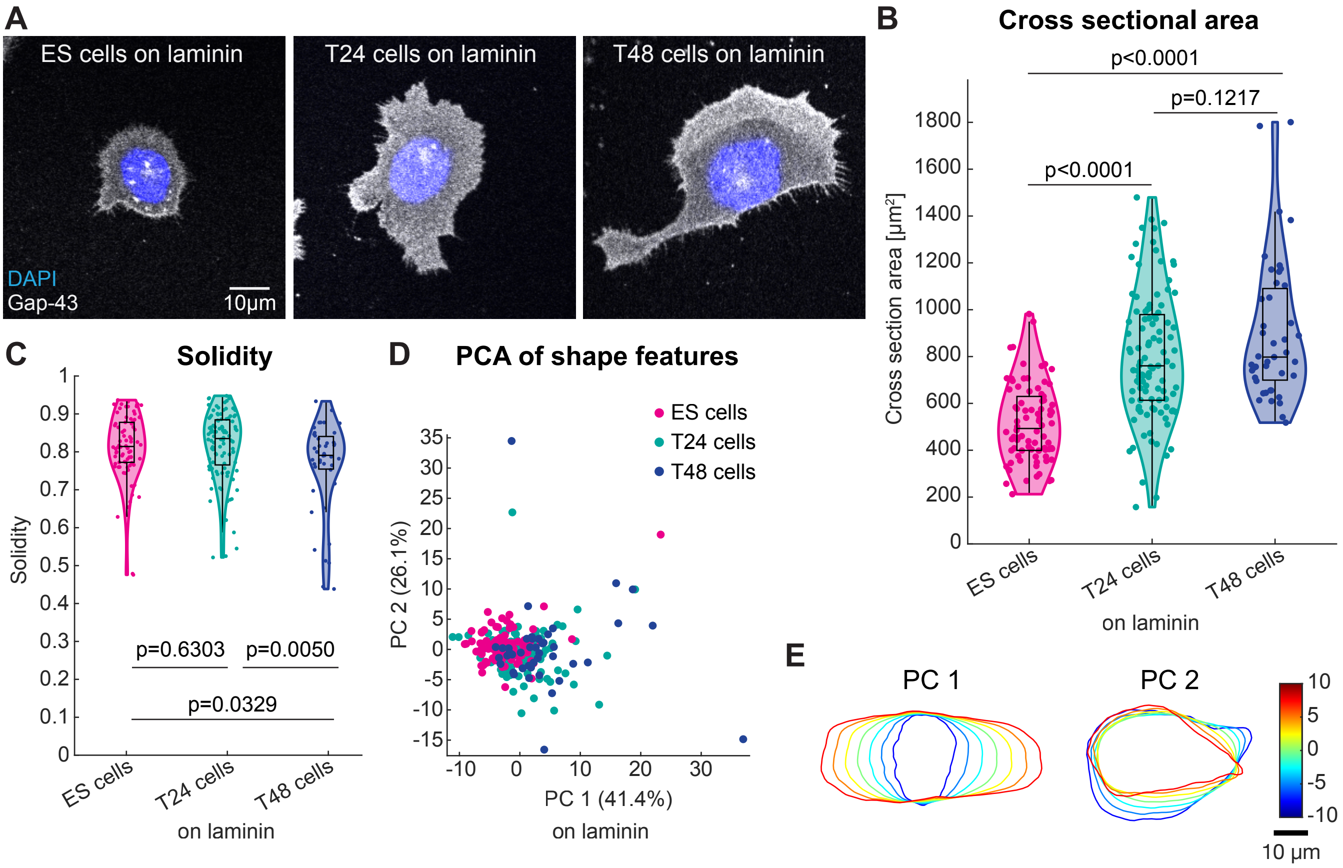

F

| Cells | Cross sectional area | Solidity | n | N |
|---|---|---|---|---|
| ES | 493 ± 164 μm² | 0.814 ± 0.088 | 82 | 3 |
| T24 | 760 ± 273 μm² | 0.835 ± 0.090 | 104 | 3 |
| T48 | 798 ± 301 μm² | 0.791 ± 0.123 | 43 | 3 |

### Supplementary Figure 3

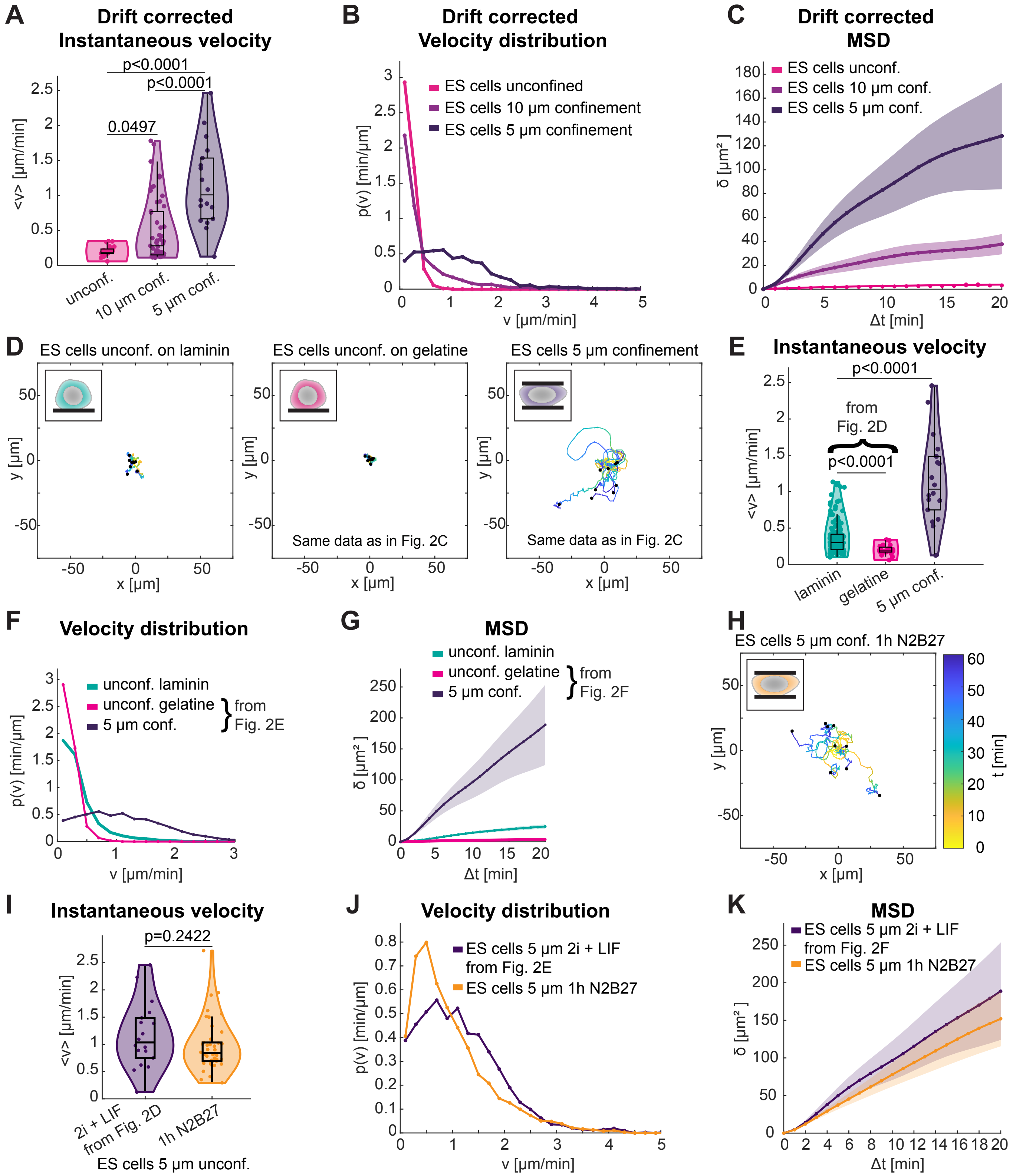

### Supplementary Figure 4

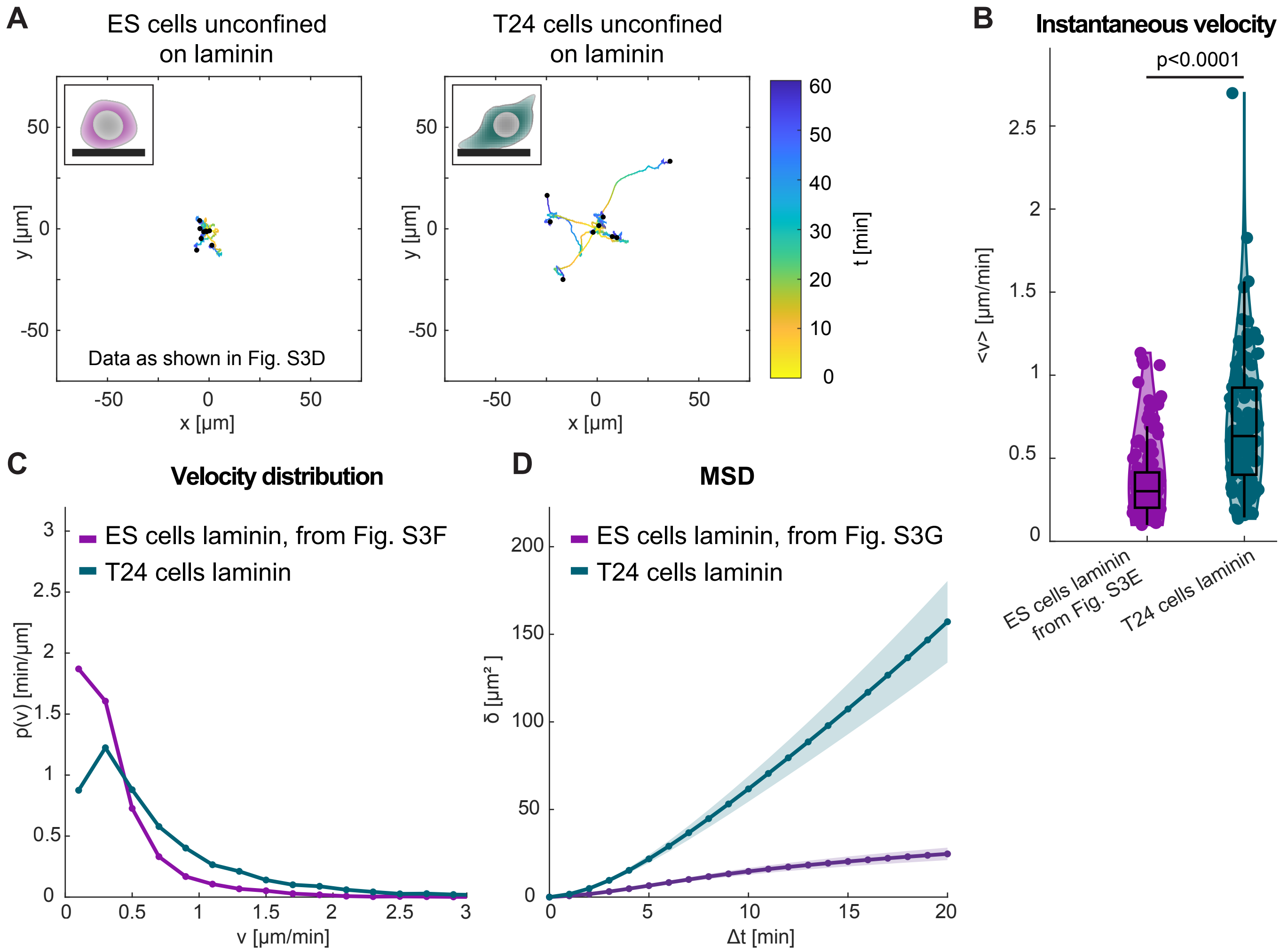

### Supplementary Figure 5

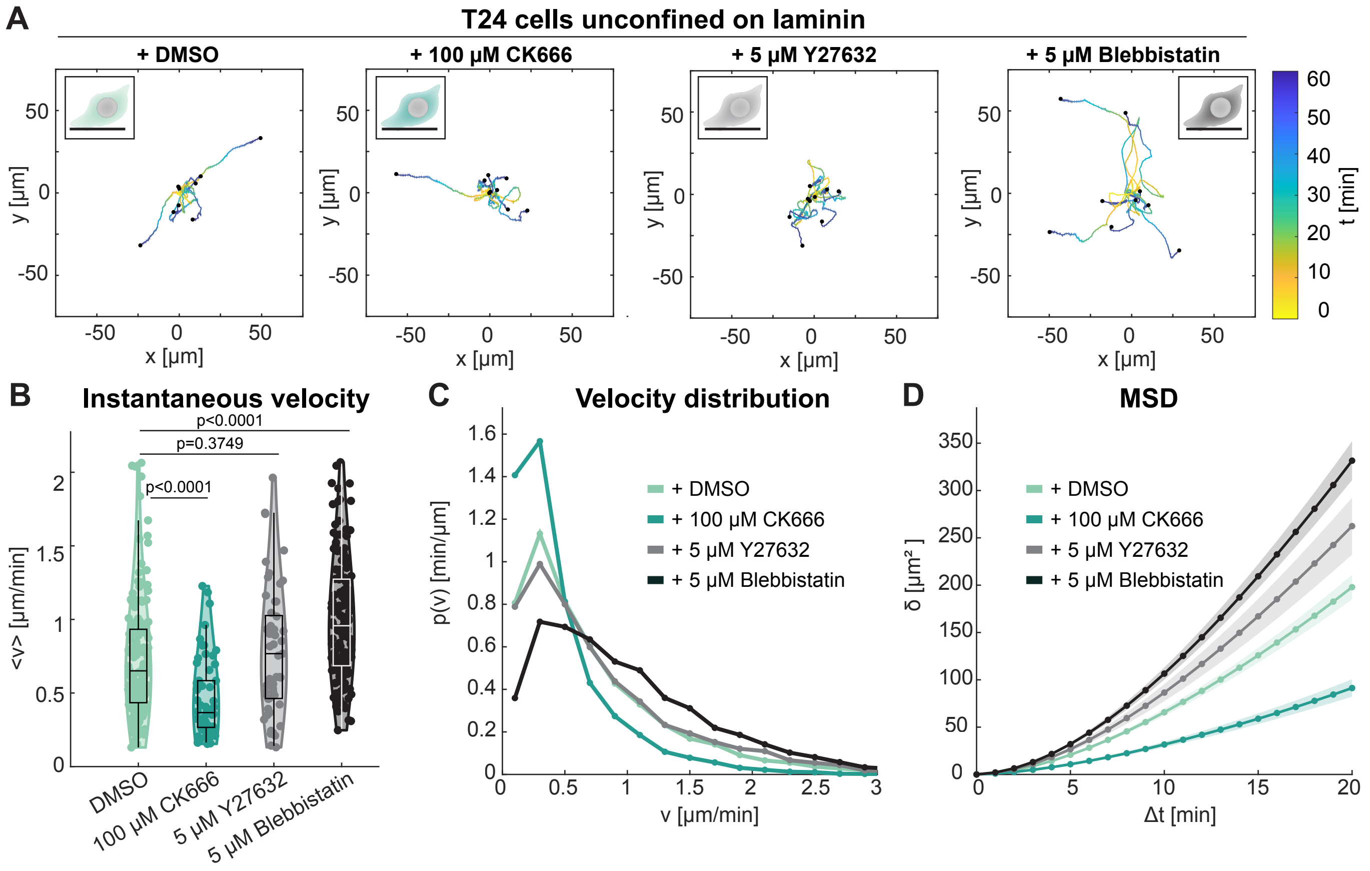

**Supplementary Table 1**

**A**

| Ribosome-depleted (total) RNA, RPKM counts (Kalkan et al., 2017) | | | | |
|---|---|---|---|---|
| **Ensembl ID** | **Gene** | **ES cells** | **T16 cells** | **T25 cells** |
| ENSMUSG00000020900 | Myosin IIA | 13923 | 12266 | 9896 |
| ENSMUSG00000022443 | Myosin IIB | 4817 | 4610 | 5781 |
| ENSMUSG00000030739 | Myosin IIC | 68 | 70 | 45 |
| **Ensembl ID** | **Gene** | **ES cells** | **T16 cells** | **T25 cells** |
| ENSMUSG00000029621 | Arpc1a | 1032 | 931 | 838 |
| ENSMUSG00000029622 | Arpc1b | 1801 | 1203 | 1305 |
| ENSMUSG00000006304 | Arpc2 | 5415 | 4540 | 3998 |
| ENSMUSG00000029465 | Arpc3 | 3228 | 2437 | 2350 |
| ENSMUSG00000079426 | Arpc4 | 3854 | 2665 | 2354 |
| ENSMUSG00000008475 | Arpc5 | 2013 | 1496 | 1524 |
